# HLA-Glyco: A large-scale interrogation of the glycosylated immunopeptidome

**DOI:** 10.1101/2022.12.05.519200

**Authors:** Georges Bedran, Daniel A. Polasky, Yi Hsiao, Fengchao Yu, Felipe da Veiga Leprevost, Javier A. Alfaro, Marcin Cieslik, Alexey I. Nesvizhskii

## Abstract

MHC-associated peptides (MAPs) bearing post-translational modifications (PTMs) have raised intriguing questions regarding their attractiveness for targeted therapies. Here, we developed a novel computational glyco-immunopeptidomics workflow that integrates the ultrafast glycopeptide search of MSFragger with a glycopeptide-focused false discovery rate (FDR) control. We performed a harmonized analysis of 8 large-scale publicly available studies and found that glycosylated MAPs are predominantly presented by the MHC class II. We created HLA-Glyco, a resource containing over 3,400 human leukocyte antigen (HLA) class II N-glycopeptides from 1,049 distinct protein glycosylation sites. Our comprehensive resource reveals high levels of truncated glycans, conserved HLA-binding cores, and differences in glycosylation positional specificity between classical HLA class II allele groups. To support the nascent field of glyco-immunopeptidomics, we include the optimized workflow in the FragPipe suite and provide HLA-Glyco as a free web resource.

## Introduction

Protein glycosylation has been extensively studied and found to play a variety of biological roles, including antigen recognition, host-pathogen interactions, and immune modulation^1^. Glycosylation causes dramatic alterations in response to cancer and has been suggested as a potential biomarker^2–5^. Moreover, glycosylation could be an attractive source of tumor-specific antigens, considering the viability of post-translational modifications (PTMs) on MHC-associated peptides^6–9^ (MAPs). Critically, glycosylation has been reported to have a significant impact on the immunogenic properties of MAPs in terms of T-cell recognition^10–12^ and epitope generation due to interference with the proteolytic cleavage^13^.

High-throughput identification of glycosylated MAPs from mass spectrometry (MS) data involves combining two notoriously challenging problems in computational proteomics. First, the proteolytic processing of MAPs requires non-enzymatic searches (*i*.*e*., non-specific cleavage of proteins at every peptide bond). Considering all possible cleavages of reference proteins results in an enormous search space of candidate sequences. Second, the non-templated nature of the glycosylation process results in hundreds of distinct glycans that can be detected across the proteome^14^. A combinatorial explosion thus takes place when considering all possible non-enzymatic peptide sequences with many possible glycans. As a result, a non-specific glycopeptide search is not feasible with many search engines due to prohibitively long run times and/or insufficient sensitivity. To the best of our knowledge, very few glycosylation analyses of MAPs have been performed. One of the earliest successful identifications of glycosylated class II MAPs was made in 2005^15^ with 2 N-linked glycopeptides found in an EBV-transformed human B-lymphoblastoid cell line. In 2017, Malaker *et al*. successfully identified 26 glycosites in 3 different melanoma cell lines^9^. Both studies required identification of glycopeptides by manual annotation of the spectra. More recently, a third effort from 2021 captured 209 unique human leukocyte antigen (HLA) II-bound peptide sequences from the SARS-CoV-2 virus^16^ using an automated glycopeptide search method assisted with a manual verification of all glycopeptide spectra.

Large-scale analysis of glycosylated MAPs requires automated methods with exceptional speed and accuracy to handle the enormous search space of glycosylated non-specific peptides. The above-mentioned challenges have been tackled by our recent developments to improve the search speed^17^ (MSFragger) and address the complexity of glycosylation^18^ (MSFragger-Glyco). Building on these advances, we developed an optimized workflow for HLA glyco searches with a focus on optimizing the false discovery rate (FDR) control of glycosylated MAPs. We assembled, carefully annotated, and analyzed 8 publicly available immunopeptidomic datasets for N-glycosylation using our workflow and investigated the glycosylated MAPs binding properties. From nearly 2,000 LC-MS/MS runs, we found 3409 class II N-glycosylated MAPs on 1049 distinct protein glycosylation sites of 677 unique proteins. We revealed characteristics of HLA glycopeptides, including high levels of truncated glycans, conserved HLA-binding cores across the 72 studied HLA class II alleles, and a different glycosylation positional specificity between the classical allele groups.

Induced expression and antigen-presentation by the MHC class II on tumor cells is increasingly being recognized as a mediator of anti-tumor immunity and neoantigen efficacy^19–24^. Our results, made readily accessible as a free web resource, significantly expand our understanding of glyco-MAPs in cancer; and together with our novel optimized workflow, are expected to further the development and discoveries in the nascent field of glyco-immunopeptidomics.

## Results

### Computational glyco-immunopeptidomics workflow

The computational workflow developed in this work for the analysis of glycosylated MAPs is illustrated in **Fig. 1**. While O-glycosylated MAPs are also of potential interest^25^, O-glycopeptide analysis typically requires electron-based activation to locate the glycosite(s) within the peptide. As the vast majority of available immunopeptidomics data lacks such activation, we focused exclusively on N-glycosylated MAPs for this analysis. Briefly, MSFragger-Glyco performs N-glycosylation motif checks for the N-X-S/T consensus sequence, which serves as the attachment site for polysaccharides (*i*.*e*., sequon). Simultaneously, spectra are checked for the presence of fragmentation products of peptide-conjugated glycans (*i*.*e*., oxonium ions). The glycan search is only performed for peptides with a sequon and for spectra containing oxonium ions above a relative intensity threshold (10% in this case). A regular search is performed for all other spectra. Next, we use PeptideProphet^26^ and ProteinProphet^27^ within the Philosopher^28^ toolkit to model and filter false discovery rates (FDR) to 1% for peptide-spectrum matches (PSMs), peptides, and proteins, respectively. As in previous glycopeptide analyses, we applied the extended mass model of PeptideProphet to simultaneously model the score and mass-shift distributions of the database search^17^. This provides a separate probability model for different glycan masses (i.e., mass shifts) to account for the varying frequencies of the different glycans^18^.

**Figure 1:**
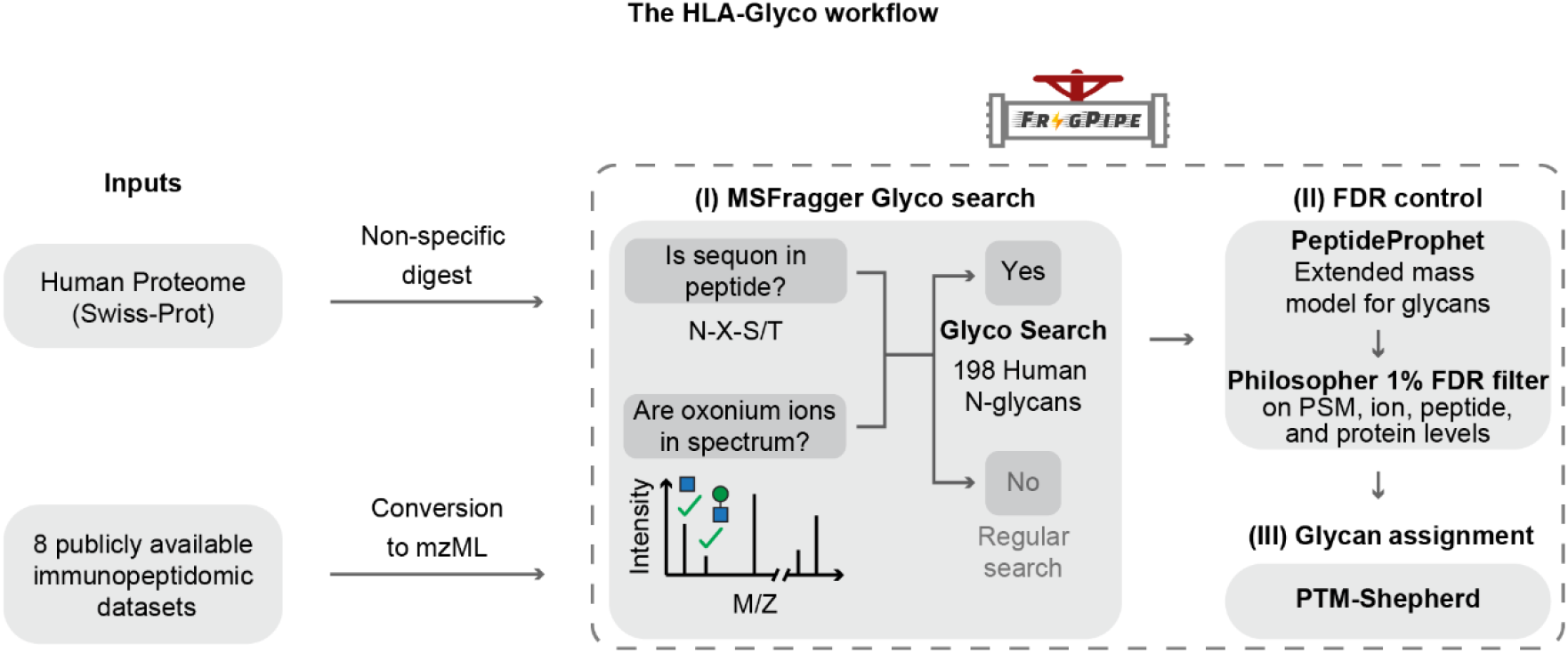
The HLA-Glyco workflow for the detection of glycosylated MHC associated peptides. The FragPipe suite was used to (I) perform a search for glycosylated peptides (glyco search) with the MSFragger search engine; (II) control the FDR with PeptideProphet in combination with a modified version of Philosopher; and (III) assign a glycan composition for each glycopeptide-spectrum match using PTM-shepherd.

Initially, we assessed the standard FDR procedures used for enzymatically digested and enriched glycopeptides on non-enzymatic unenriched immuno-glycopeptides. We observed that while 91% of the glycoPSMs corresponded to known glycosylation sites, less than half of the observed glycosites (46%) were previously known (**Supplementary Fig. 1a**). Thus, known sites tended to have many supporting spectra, while unknown sites had few and notably lower scores, likely indicating an unacceptable increase in false discoveries. Since glycoPSMs represent a small fraction of the identified spectra, the score thresholds used in our initial FDR filtering were mostly influenced by non-glycosylated peptides. As glycopeptides have a much larger search space, this results in an enrichment of false discoveries in the glycopeptide fraction when all PSMs are filtered together. To counter this, we applied a separate PeptideProphet probability (*i*.*e*., score) filter for glycosylated and non-glycosylated PSMs to control FDR in each category despite the differences in search space, using a modified version of Philosopher (see **Methods** and **Supplementary Fig. 1b**). We further filtered glyco-PSMs by glycan q-value (q ≤ 0.05) to remove glycopeptides lacking sufficient evidence supporting the glycan composition assignment^29^ by PTM-Shepherd^30^. With this improved filtering method, the proportion of PSMs corresponding to known glycosites increased to 96%, and the proportion of identified glycosites corresponding to known glycoproteins increased to 95%, with 79% of sites previously identified in other glycoproteomic analyses **(Supplementary Fig. 1c**). These stringent glycopeptide-specific filters provide effective FDR control in a challenging search, allowing for confident construction of the HLA glycopeptide resources.

### Large multi-tissue MHC immunopeptidome dataset

We selected 8 immunopeptidomic studies^31–38^, prioritizing studies with a large amount of high-resolution mass spectrometry data and included a variety of instruments as a means to reduce instrumental bias (see **Methods**). Based on our careful curation and annotation of these data, our collection of 732 different HLA class II mass spectrometry samples incorporated 90.8% of HLA-typed data (**Fig. 2a**), 80.3% of patient tissues, 16.7% of cell lines, and 2.9% of tumor-infiltrating lymphocytes (**Fig. 2b**). The previously mentioned sample types covered up to 6 different cancers (**Fig. 2c**) located in the brain (meningioma and glioblastoma), skin (melanoma), colon (colorectal), and lung (adenocarcinoma and squamous carcinoma). In addition, 59% of the samples are non-cancerous and come from disease-free individuals. In terms of HLA diversity, up to 72 HLA class II alleles of the 3 classic genes (DP, DQ, and DR) are covered by varying numbers of mass spectrometry samples (**Fig. 2d**).

**Figure 2:**
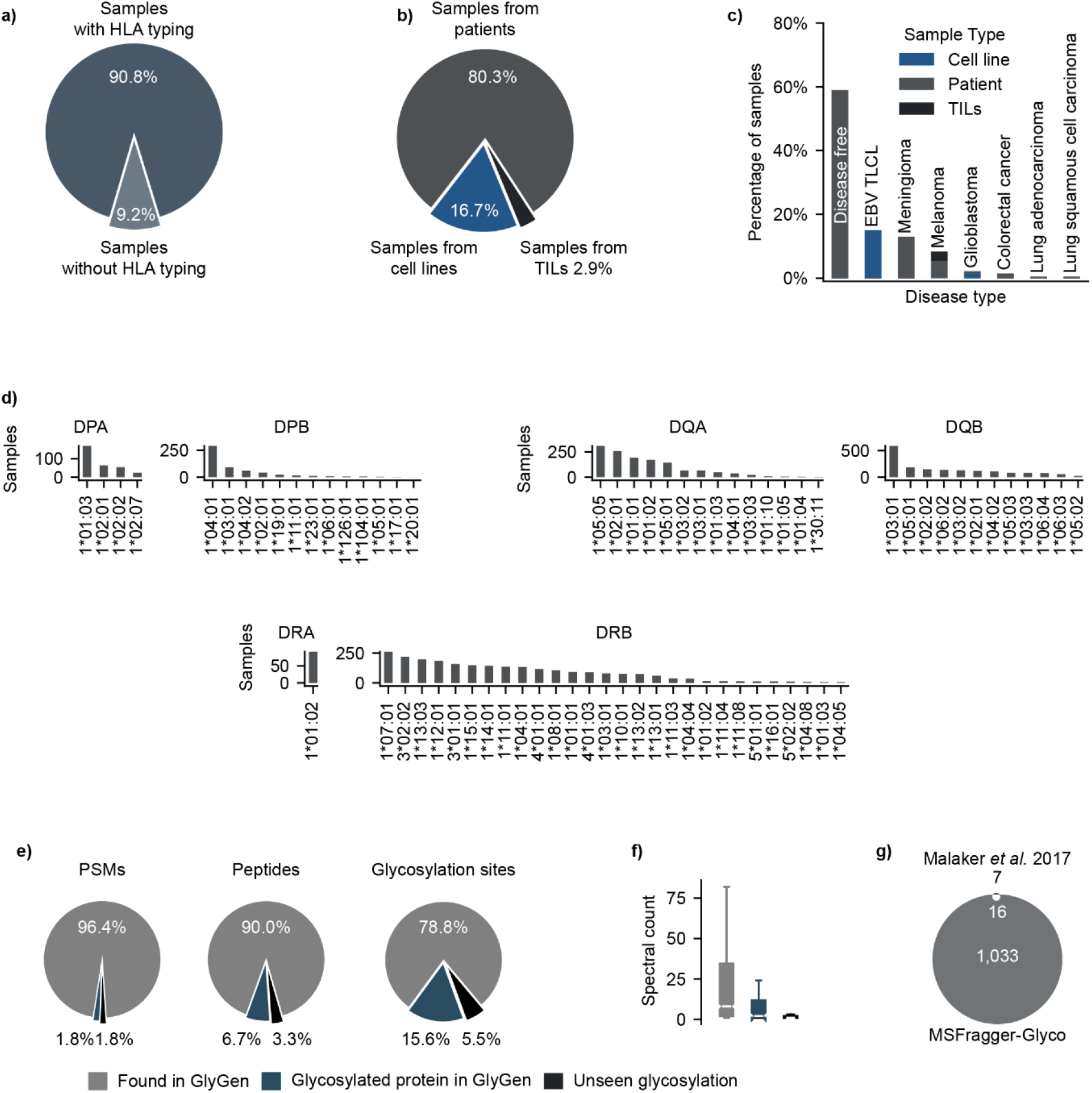
HLA class II infographics of the 8 collected datasets in this study. **a)** Percentage of samples with HLA class II typing information. **b)** Sample types of the collected mass spectrometry samples (i.e., patient tissues, cell lines, and tumor-infiltrating lymphocytes/TILs). **c)** Cancer types across the collected mass spectrometry samples. **d)** HLA class II alleles (DR, DB, and DQ) across the collected mass spectrometry samples. **e)** Percentage of glyco-PSMs, glycopeptides, and glycosylation sites found in GlyGen. **f)** Abundance of the 3 categories from panel (a) by spectral count. **g)** Comparison of the identified glycosylation sites with Malaker *et al*. 2017 findings.

Leveraging the wealth of proteomic data, we queried the glycosites identified in our study against previously reported glycosylation sites in GlyGen^39^. PSM level information showed 96.4% of previously reported glycosylation sites (**Fig. 2e**), 1.8% of glycosylation sites within previously reported glycosylated proteins, and 1.8% of new glycosylation sites. On the other hand, at the peptide level, 90% of glycopeptides mapped to previously reported glycosylation sites, 6.7% of glycopeptides were within previously reported glycosylated proteins, and 3.3% contained new glycosylation sites. A similar trend was observed at the glycosylation site level, with 78.8% of previously reported glycosylation sites, 15.6% of glycosylation sites within previously reported glycosylated proteins, and 5.5% of new glycosylation sites. It appears that peptides containing previously reported glycosylation sites are abundant species, considering the high spectral count (**Fig. 2f** in gray) in comparison with the previously unreported ones (**Fig. 2f** in blue and black). We then benchmarked our findings against previous work by Malaker *et al*. 2017^9^ on glycosylated MAPs in 3 melanoma and 1 EBV-transformed B-cell lines. The original manuscript reported 93 glycosylated peptides corresponding to 26 glycosylation sites, split between N-glycosylation (23) and O-glycosylation (3). Our workflow recovered 20 of the 23 identified N-glycosylation sites, of which 4 did not pass the FDR filter. With a 45-fold increase in glycosylation sites, we identified 1033 new sites (see **Fig. 2g**).

### Enrichment of N-glycosylation in the class II immunopeptidome

Several of the datasets we searched contained both HLA class I and II peptides from the same samples and, in one case, whole proteome data, allowing us to compare the frequency and characteristics of glycosylation across these categories. Fragmentation of glycopeptides by tandem MS (MS/MS) produces highly abundant oxonium ions resulting from the fragmentation of conjugated glycan(s), which can provide an estimate of the fraction of glycopeptides in a sample prior to a database search. To understand the abundance of glycosylation at different molecular levels, we compared the percentage of oxonium-containing MS/MS scans for the 4 datasets containing multiple HLA classes (**Fig. 3a**). Interestingly, datasets A^31^ (Bassani-Sternberg *et al*. 2016), B^34^ (Chong *et al*. 2020), and D^37^ (Forlani *et al*. 2021) showed, on average, an approximate 5-fold enrichment in potential HLA class II glycosylation events compared with HLA class I data. In dataset C^32^ (Marcu *et al*. 2021), the only dataset containing samples derived from healthy tissue, a similar proportion of oxonium-containing scans was observed in the HLA class II data as in the other datasets, but there were essentially no oxonium-containing scans in the HLA class I data. As expected, the percentage of glycosylated PSMs obtained from database searches of these datasets followed a similar trend, with 0.5 to 3% of observed PSMs glycosylated in HLA class II data versus less than 0.1% glycosylated in HLA class I data (datasets A, B, and C). Strikingly, glycosylated PSMs were also enriched approximately 7-fold in HLA class II compared with the whole proteome data in dataset D (**Fig. 3b)**, a dramatic increase given the abundance of glycosylation in the proteome.

**Figure 3:**
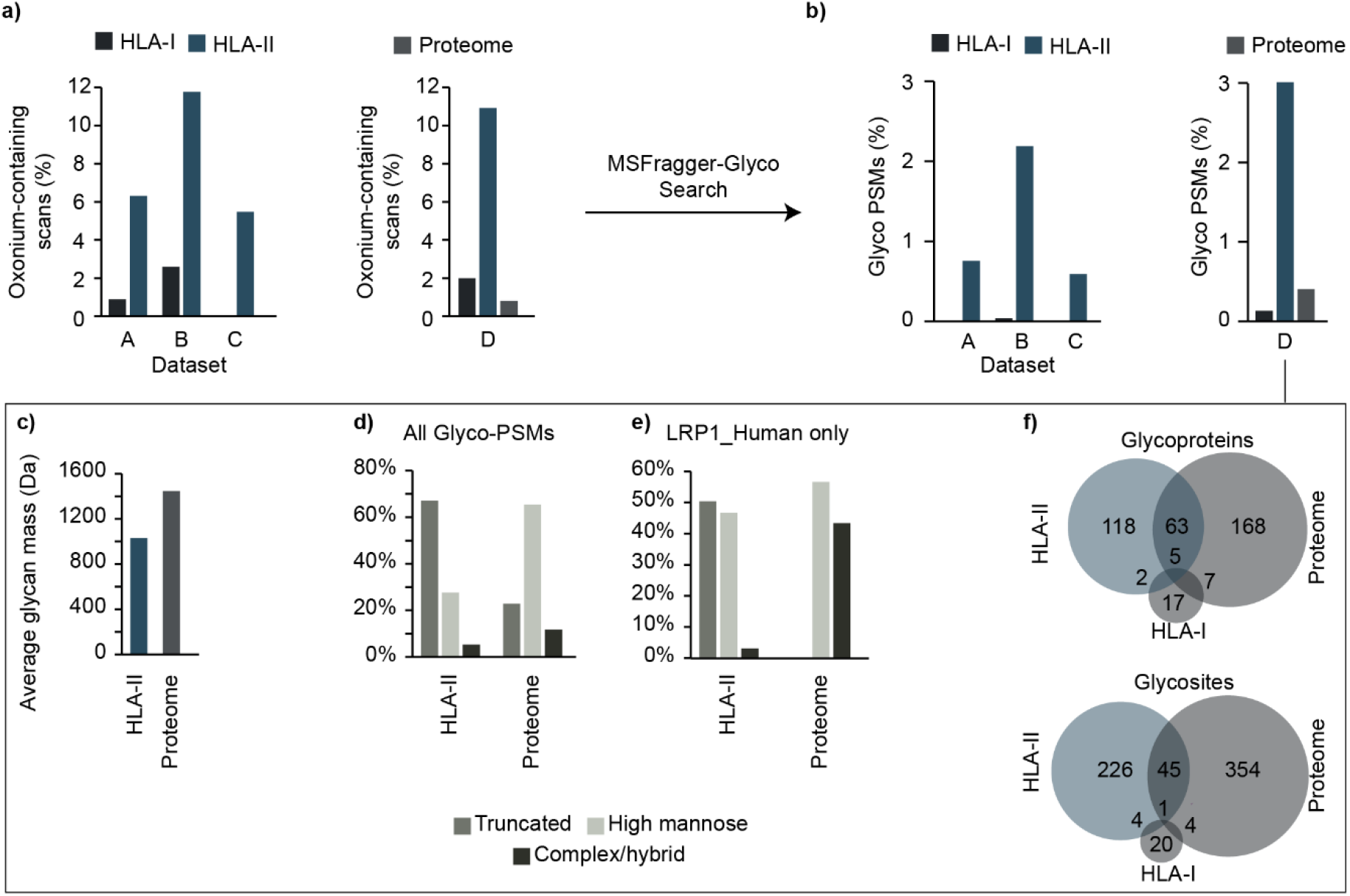
A comparison of the glycosylation on the proteome, HLA I, and HLA II peptidome levels. **a)** Levels of oxonium ions for HLA class I and II in 3 datasets (A: Bassani-Sternberg *et al*. 2016, B: Chong *et al*. 2020, C: Marcu *et al*. 2021), along with the whole proteome in dataset D: Forlani *et al*. 2021. **b)** Percentage of Glycosylated PSMs for the HLA class I and II immunopeptidome in 3 datasets (A, B, C), along with the whole proteome in dataset D. **c)** Average glycan mass in Dalton (Da) for the HLA class II immunopeptidome versus the whole proteome in dataset D. **d)** Glycan types for the class II immunopeptidome versus whole proteome in dataset D. **e)** Glycan types found in the low-density lipoprotein receptor-related protein 1 (LRP1) for the class II immunopeptidome versus the whole proteome in dataset D. **f)** Comparison of glycoproteins (top) and glycosites (bottom) found in the HLA class I, II immunopeptidome, and whole proteome of dataset D.

We also noticed that the composition of glycans observed in the immunopeptidomic datasets was different from that of their proteome counterparts. (**Fig. 3c)**. The average glycan mass detected in the immunopeptidome was approximately 1000 Da, which was significantly lower than that observed in the proteome (1400 Da average). To further explore the nature of this compositional discrepancy, we compared glycan types between the two groups (**Fig. 3d**). A higher percentage of truncated glycans (68%) was observed in the HLA class II immunopeptidome compared to the more typical high-mannose and complex/hybrid categories in the proteome, as noted in a previous analysis^9^. This trend of truncated glycans on HLA peptides was preserved when only glycans from the same protein were considered. For example, LRP1, a highly glycosylated protein, was observed with a mix of high-mannose and complex glycans in the proteome sample, but with a mix of truncated and high-mannose glycans in the HLA-II sample with almost no mature complex glycans detected (**Fig. 3e)**. There was very little overlap between the glycosylated proteins and sites in each category, with only 22.8% of HLA-II glycoproteins observed in the whole proteome data and even lower overlap (16.3%) when considering the specific glycosylation sites within proteins. (**Fig. 3f**). The whole proteome glyco search likely captures glycopeptides from the most abundant glycoproteins, as the experiment was performed without any glycopeptide enrichment, whereas the immunopeptide datasets presumably capture MAPs with much less dependence on overall protein abundance.

Overall, the data showed a remarkable enrichment of glycosylation in HLA class II-associated peptides relative to HLA class I and the whole proteome, leading us to focus the remainder of our efforts on HLA class II-associated and glycosylated peptides.

### Glycosylation of MAPs does not influence the HLA binding motif

To explore glycosylation in the context of HLA class II presentation, we focused on the HLA-binding core, a 9-mer sequence that interacts with the HLA molecule. In most mass spectrometry experiments, samples express multiple HLA alleles, leading to an ambiguous association between the identified peptides and the pool of available HLA molecules. Hence, a deconvolution step to find the HLA motifs and the corresponding binding core offsets of each peptide was deemed necessary for further experimentation (see **Methods**).

### Deconvolution of peptides using a semi-supervised approach

We first chose to use MoDec^38^ for deconvolution, a fully probabilistic framework that learns both the motifs and preferred binding core position offsets from the sequences themselves. The fact that MoDec does not rely on a pre-trained model is crucial when exploring HLA-bound peptides with post-translational modifications (*i*.*e*., glycosylation) to avoid the removal of all peptides that were not well modeled. Such a deconvolution strategy requires manual intervention to choose the number of HLA motifs (*i*.*e*., number of clusters) and assign each discovered motif to one of the expressed HLA alleles of a given sample. We carefully selected a case study on a human B lymphoblastoid cell line (C1R) from Ramarathinam *et al*. 2021^36^. The purification protocol of the HLA-bound peptides in this study was performed sequentially with pan anti-class I, followed by class II anti-DP (**Fig. 4a**), class II anti-DQ (**Fig. 4b**), and class II anti-DR antibodies (**Fig. 4c and d)**. Hence, the resulting mass spectrometry samples were mono-allelic (*i*.*e*., presenting one allele at a time), except for the DR samples with the DRB1*12:01 and DRB3*02:02 alleles eluting together. **Fig. 4** presents 4 sections **a, b, c**, and **d** standing for the HLA class II alleles DPA1*02:01/02-DPB1*04:01, DQA1*05:05-DQB1*03:01, DRB1*12:01, and DRB3*02:02, respectively. All alleles showed a similar percentage of glycosylated and non-glycosylated peptides with the corresponding HLA motifs after deconvolution (**Fig. 4, panel I**). All 25 replicates showed an unaltered HLA-binding core with glycosylation (two-sided Fisher’s exact test, 25 P-values > 0.05).

**Figure 4:**
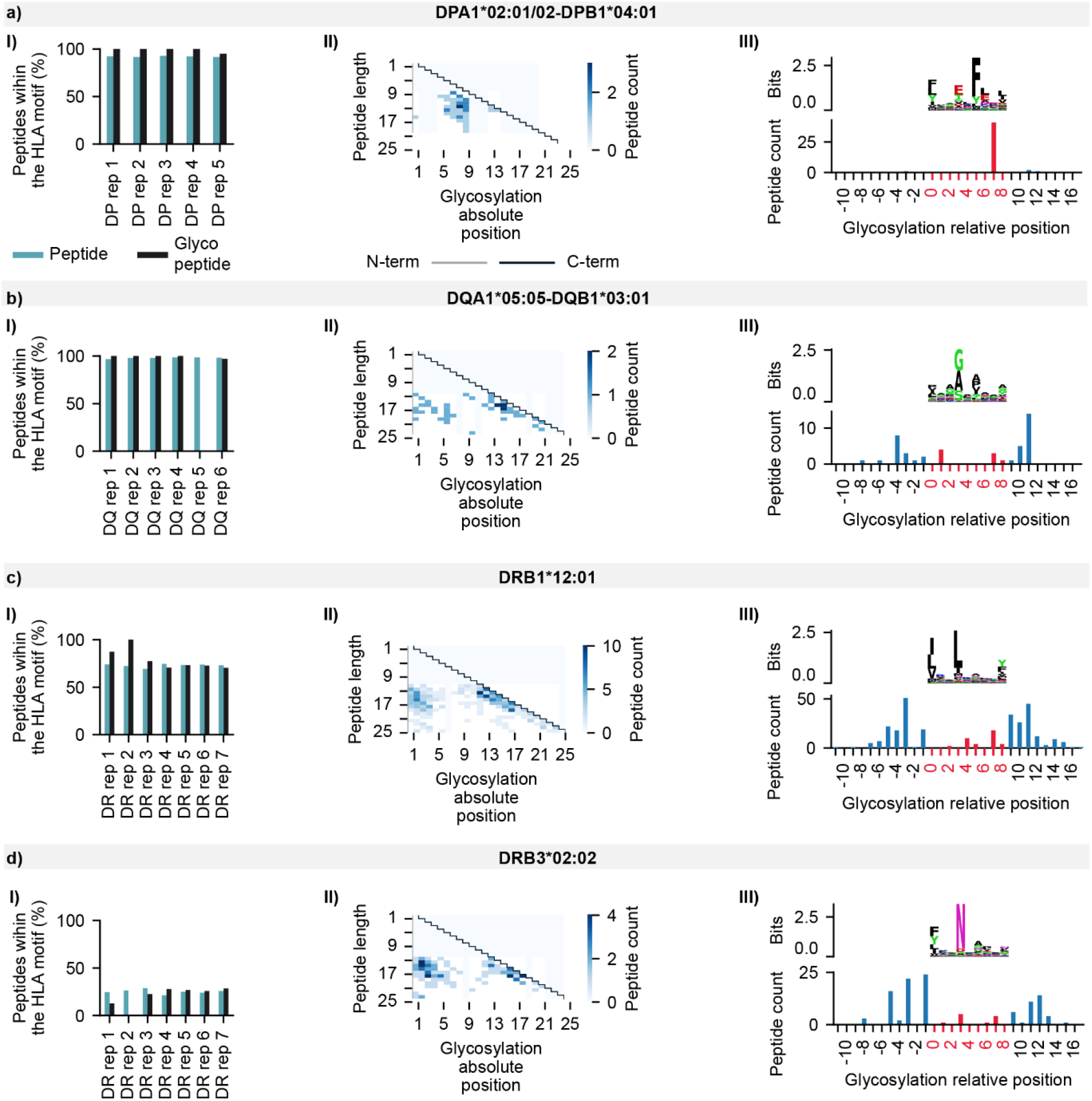
Semi-supervised deconvolution of glycosylated HLA peptides from Ramarathinam *et al*. 2021 using MoDec. **Panels I** show the percentage of peptides and glycopeptides presenting the HLA binding motif. **Panels II** display the glycosylation absolute position within the peptidic sequence (x-axis) and the peptide length (y-axis). Gray and black lines indicate the N-term and C-term respectively while the white to blue gradient represents the number of peptides with a specific glycosylation position at a specific peptide length. **Panels III** present the HLA binding motif after deconvolution with MODEC (top) and the number of glycopeptides per relative glycosylation position (bottom). Negative values refer to glycosylation position upstream the HLA-binding core, values between 0 and 8 represent positions within the HLA-binding core, and values ≥ 9 refer to positions downstream the HLA-binding core. **a)** Peptides associated with the HLA allele DPA1*02:01/02-DPB1*04:01 of the C1R cell line. **b)** Peptides associated with the HLA allele DQA1*05:05-DQB1*03:01 of the C1R cell line. **c)** Peptides associated with the HLA allele DRB1*12:01 of the C1R cell line. **d)** Peptides associated with the HLA allele DRB3*02:02 of the C1R cell line.

Considering the concordance of glycopeptide sequences with the HLA-binding cores, we checked the absolute glycosylation position per peptide length (*i*.*e*., glycosylation offset within the peptide). **Fig. 4** panel **II** shows a glycosylation tendency towards the N- and C-termini for both DQ and DR alleles (**Fig. 4** sections **b, c**, and **d** at panel **II**) and only the C-terminal tendency for the DP allele (**Fig. 4** section **a** at panel **II**). To further decipher glycosylation in the context of the HLA-binding cores, we looked at the relative position shown in **Fig. 4** panel **III** (*i*.*e*., glycosylation offset from the HLA-binding core start). Negative values indicate sites upstream of the HLA-binding motif start, 0 to 8 values reference positions within the HLA-binding core, and values greater than 8 denote glycosylation sites downstream of the HLA-binding core. For the DPA1*02:01/02-DPB1*04:01 allele, glycosylation occured 91% of the time within the HLA motif at position 8 (**Fig. 4** section **a** at panel **III**). In contrast, for the other 3 alleles, glycosylation was more likely (86% of the time) to take place up- or downstream of the HLA-binding core.

### Deconvolution of peptides using a fully unsupervised approach

Despite the usefulness of MoDec for a previously unexplored category of peptides, such a tool suffers from several limitations^40,41^: (I) the need for manual intervention to associate the identified motifs with known allele specificities present in the sample; (II) the difficulty of assigning peptides to MHC molecules when alleles with overlapping motifs are co-expressed; (III) low sensitivity with low expression of MHC molecules; and (IV) the complexity of HLA class II specificities due to the involvement of the variable alpha and beta chains for the HLA-DQ and HLA-DP groups. All these, render motif-allele assignment a daunting task, especially with up to 87 subjects in our dataset. Thus, we used the state-of-the-art binding model NetMHCIIpan 4.1^41,42^ to perform MHC motif deconvolution and assign glycopeptide sequences to their most likely HLA alleles without the need for manual intervention (see **Methods**). Consistently, glycosylated and non-glycosylated peptides from Ramarathinam *et al*. 2021 showed similar binding properties, indicating that the detected glycosylation fit within the known HLA-binding cores (two-tailed Fisher’s exact test, P-value: 0.48). Interestingly, NetMHCIIpan 4.1 confirmed most peptides with glycosylation located at P8 within the HLA-binding core (97% for DPA1*0201 and 100% for DPA1*0202) for the C1R DP allele (**Fig. 5a**). Overall, 95%, 83%, 76%, and 87% of glycopeptides were found to bind to C1R DP (**Fig. 5a**), DQ (**Fig. 5b**), DRB1*12:01 (**Fig. 5c**), and DRB3*02:02 (**Fig.5d**), respectively. Hence, we carried out the NetMHCIIpan 4.1 deconvolution for the 83 remaining subjects in our dataset.

**Figure 5:**
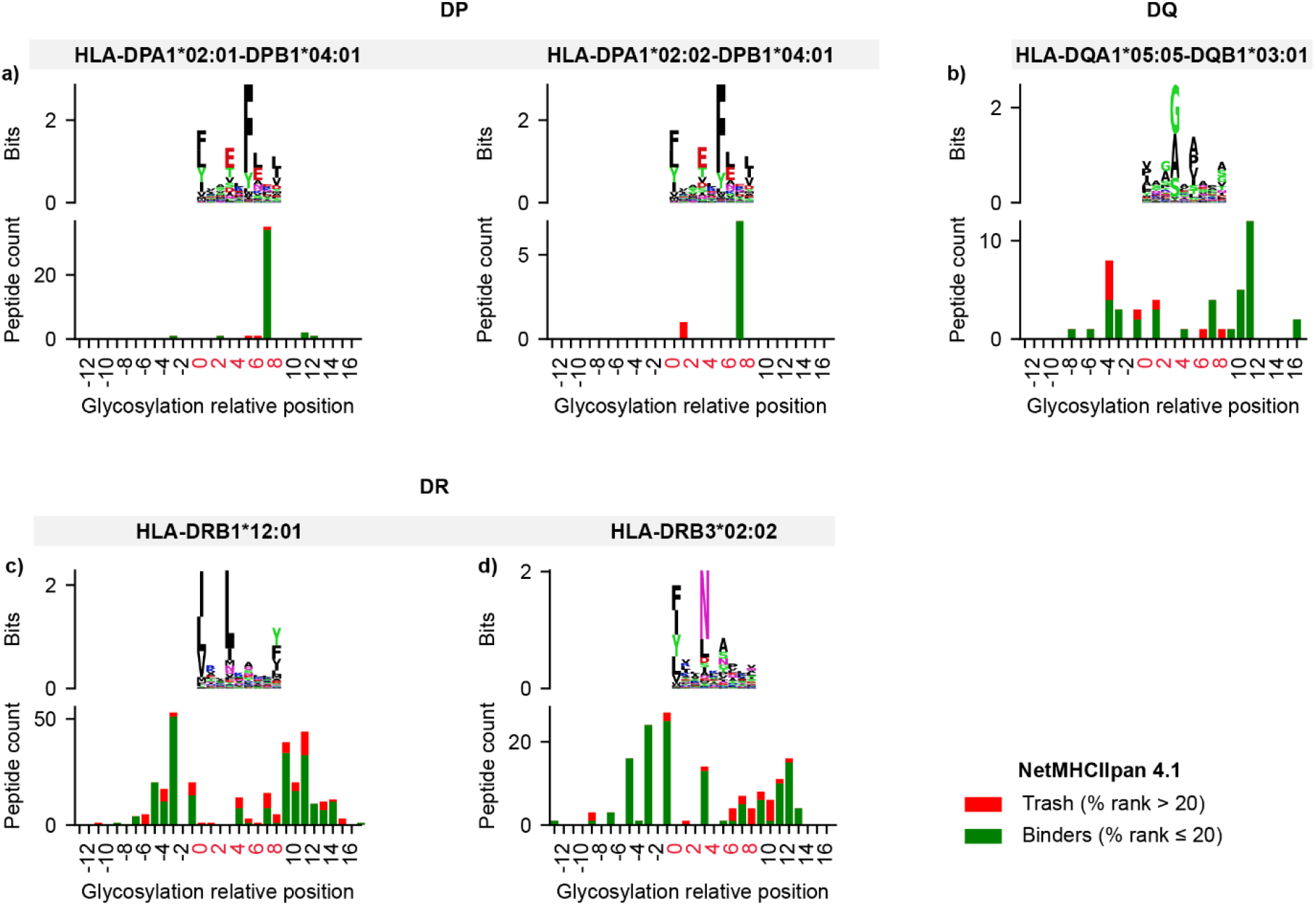
Fully unsupervised deconvolution of glycosylated HLA peptides from Ramarathinam *et al*. 2021 with NetMHCIIpan 4.1. Each panel illustrates 2 levels of information: the top level shows the HLA-binding motif of peptides passing a NetMHCIIpan 4.1 percentile rank threshold of 20 after binding affinity prediction. The bottom level shows glycopeptides that are predicted to bind to a given allele in green (%rank ≤ 20), otherwise non-binder peptides (*i*.*e*., trash) are shown in red (%rank > 20). Positions are shown relatively to the HLA binding core with negative values referring to glycosylation position upstream the HLA-binding core, values between 0 and 8 represent positions within the HLA-binding core, and values ≥ 9 refer to positions downstream the HLA-binding core. **a)** Deconvolution of glycosylated peptides associated with the HLA-DPA1*02:01/02-DPB1*04:01 alleles. **b)** Deconvolution of glycosylated peptides associated with the HLA-DQA1*05:05-DQB1*03:01 alleles. **c)** Deconvolution of glycosylated peptides associated with the HLA-DRB1*12:01 allele. **d)** Deconvolution of glycosylated peptides associated with the HLA-DRB3*02:02 allele.

### The HLA class II N-glycosylation characteristics

We noticed a high tendency of glycosylation within the HLA-binding core for HLA DP alleles, followed by a lower tendency for HLA DQ, and even lower one for HLA DR alleles. Hence, we checked for the occurrence of such events for each of the 3 HLA groups (DP, DQ, and DR). **Fig. 6a** shows that up to 57% of HLA DP associated peptides have glycosylation inside the HLA-binding core, 30% for HLA DP, and 13% for HLA DR. In terms of glycan types, **Fig. 6b** shows that HLA DP associated peptides showed the highest fraction (0.67) of truncated glycans compared to DQ (0.55) and DR (0.41). High-mannose glycans showed a reverse trend for DR, DQ, and DP alleles, with fractions of 0.37, 0.27, 0.21, respectively. All DP, DQ, and DR associated peptides showed a depletion in complex/hybrid glycans in accordance with previous findings^9,16^.

**Figure 6:**
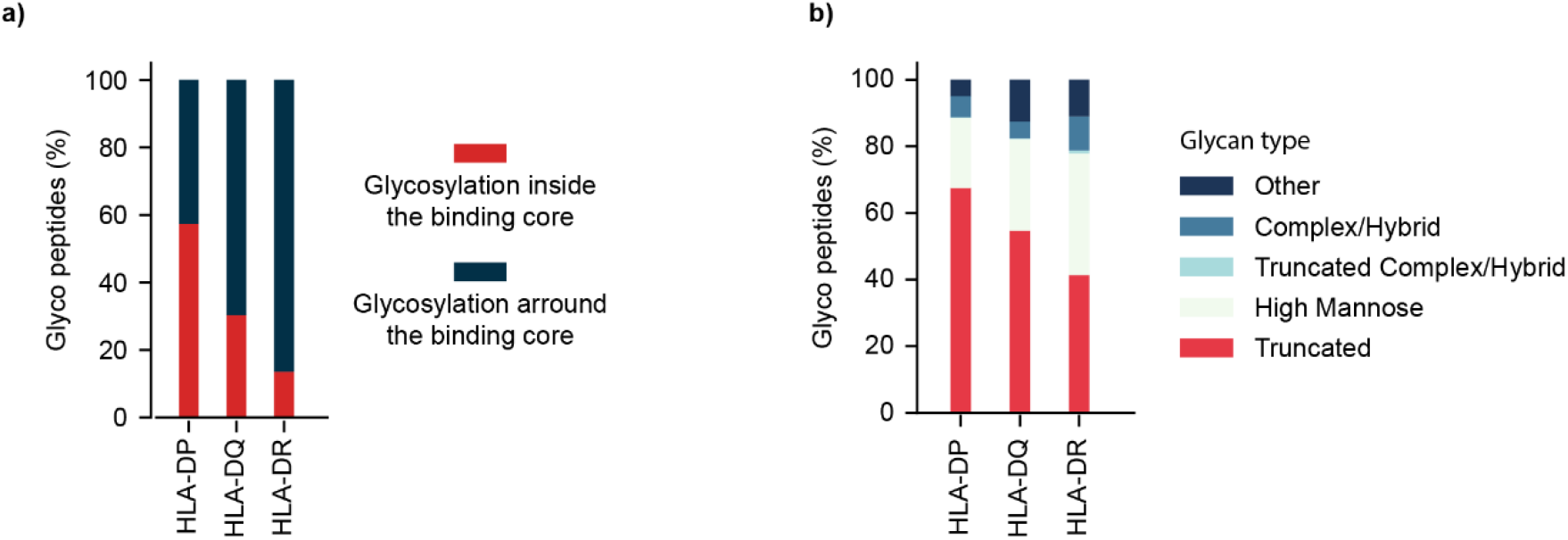
Glycan characteristics of the glycosylated HLA class II associated peptides. **a)** Percentage of glycosylation inside (red) and outside (blue) the HLA binding motif per HLA group (DP, DQ, and DR). **b)** Distribution of glycan types among the studied HLA class II group (DP, DQ, and DR).

## Discussion

Post-translational modifications increase the diversity of the immunopeptidome and may provide new targets for the immune system to recognize tumor cells or respond to pathogens. With PTM-driven antigenicity being continuously highlighted^9,31,43,44^, glycosylation is a key PTM that, despite its long history of research, remains understudied in the context of MHC presentation due to computational related challenges. In this work, we have developed a workflow for glyco-immunopeptidomics that combines the speed and sensitivity of MSFragger-Glyco, with the inclusion of glycopeptide-specific FDR control in Philosopher, which is critical for filtering out low-confidence identifications. We used this workflow to produce a resource of HLA class II N-glycosylated MAPs arising from a harmonized analysis of 8 publicly available studies. Overall, we identified 1049 glycosylation sites from 3409 different glycopeptides, an order of magnitude greater than any previous effort in this area. Leveraging this large-scale resource, we explored the properties of glycosylated MAPs, including the types of glycans conjugated, MHC binding affinity predictions, and the positioning of glycosylation relative to the HLA binding core. Interestingly, we observed no difference in binding motif predictions with glycopeptides compared to non-glycopeptides, despite some peptides containing glycans within the binding core. HLA DP alleles presented a majority of glycans within the binding core (57%) compared with HLA DQ alleles (30%) and HLA DR alleles (13%). Moreover, we found a difference in the glycan types between HLA groups (DP, DR, and DQ), with truncated glycans enriched for DP alleles and a higher mannose content for DR alleles.

A study by Malaker *et al*.^9^ on HLA class II N-glycosylation covered 5 DR alleles (DRB1*0101, DRB1*0401, DRB1*0404, DRB1*1502, DRB4*0103) and showed that 3 out of 23 peptides had glycosylated residues within the binding core. In combination with molecular modeling, this allowed the authors to postulate that glycan residues are most likely to protrude out of the HLA-binding pocket and interact with the complementary determinant region of the T-cell receptor. Our findings expand the coverage to 28 DR alleles, along with multiple DP and DQ alleles, adding up to 87 HLA molecules overall, when considering the combination of alpha and beta chains. In addition to the preference of terminal glycosylation for peptides associated with DR and DQ alleles, we observed an HLA-binding core glycosylation tendency for peptides associated with DP alleles. Future studies should explore whether the correlation between smaller glycans and presence within the HLA-binding core is related simply to size restrictions preventing larger glycans from occupying the core or is a reflection of other processing of MAPs for presentation.

The enrichment of glycosylated peptides on the MHC-II, while preserving canonical binding motifs, offers the tantalizing possibility of designing and developing glycosylated neoantigen vaccines with improved affinity over wild-type peptides^22,23^. Which is further notable, in light that most of the known anti-tumor CD4+ T cells are specific for highly immunogenic self-derived MHC-II antigens, demonstrating that self-antigen CD4+ T cells can mount anti-tumor responses. Cancer-specific glycosylation of MAPs may further contribute to the restriction of those mechanisms to the tumor microenvironment. We made our findings readily available as a web resource to query pertinent information about the identified glycosylated MAPs. Users can search for a specific glycan and/or MAP sequence, protein, or glycosylation site associated with a specific HLA allele. In addition, we included deconvolution information allowing further interpretation of the data within the HLA haplotype context. We are planning to grow this initiative, introduce more studies, and increase the HLA allele coverage. Moreover, by providing the optimized computational workflow file, which can be loaded directly into FragPipe to reproduce the method described here, we make it easy for others to carry out challenging glyco-immunopeptidomics analyses on new datasets. It is our hope that the method and findings presented here will expand the field of tumor-specific antigen discovery, broaden the scope of possible antigens to target, and improve strategies for vaccine design. O-glycosylated MAPs, for example, represent another potential class of antigens that can, in principle, be studied by our method for further exploration^45^. Finally, given the promising nature of glycosylated MAPs, we anticipate the attraction of glycosylation-oriented research towards the immunopeptidomics field.

## Methods

### Dataset selection

Studies from the PRIDE^46^ database were first screened based on a list of keywords related to immunopeptidomics. Next, low-resolution analyses were eliminated, and MHC-related datasets conducted with at least one of the following instruments were kept: Orbitrap Lumos/Fusion, Q Exactive, LTQ Orbitrap, Orbitrap Exploris 480, TripleTOF, impact II, and maXis. Then, manual curation of the resulting 312 studies was performed to filter non-relevant datasets, resulting in 140 HLA Class I, II, or I & II datasets. The number of identified proteins per study was retrieved from gpmDB^47^ and datasets with a high number of protein groups were prioritized. A final manual curation step resulted in the selection of the 8 datasets included in this study.

### Mass spectrometry N-glycan search

Raw and wiff files were first downloaded from PRIDE and converted to mzML format using msconvert^48^ with peak picking, FragPipe (TPP) compatibility, and removal of zero values filters. The analysis was executed within the FragPipe suite v18.1-build5 using headless mode. Glyco-searches were performed using MSFragger v3.5 with methionine oxidation, N-terminal acetylation, and cysteinylation as variable modifications, and a list of 198 glycans. A list of contaminants was added to the UniProt Swiss-Prot (UP000005640) proteins^49^, along with their corresponding reversed decoy sequences. Enzymatic cleavage was set to non-specific with peptide lengths from 7 to 25 amino acids for the 8 HLA class II datasets, from 7 to 12 amino acids for HLA class I datasets (A, B, C, D), and fully enzymatic cleavage with peptide lengths from 7 to 50 amino acids for the whole proteome dataset D. Peptides containing the consensus sequon (N-X-S/T) and decoy (reversed) peptides containing the reversed sequon were considered as potential glycopeptides to ensure the that same number of potential glycopeptides was searched in both target and decoy databases. Only spectra containing oxonium ion peaks with summed intensity of at least 10% of the base peak were considered for glycan searches, while all others were searched without considering glycosylation. Data were deisotoped^50^ and decharged in MSFragger-Glyco, calibrated, and searched with 20 ppm mass tolerances for precursors and 15 ppm for products with MSFragger’s built-in parameter optimization performed for each study^51^. Errors in monoisotopic peak detection by the instrument were allowed (+1 and +2 Da).

### FDR control

Filtering was performed using Philosopher^28^ (v4.5.1-RC10), including PeptideProphet modeling of peptide probabilities, ProteinProphet protein inference, and Philosopher’s internal filter for FDR control. The semi-parametric modeling of PeptideProphet was used with the expectation value as the only contributor to the f-value. The number of tolerable termini (ntt) model was disabled, as it is not applicable to non-enzymatic searches. Filtering was performed in Philosopher using a modified, group-specific FDR procedure. Non-glycosylated and glycosylated PSMs were filtered separately, using a delta mass cutoff of 145 Da (the size of the smallest glycan considered in the search) to distinguish glycosylated PSMs from non-glycosylated PSMs. This allowed different score thresholds to be used to filter glycosylated and non-glycosylated PSMs to 1% FDR. This is essential as the large search space for glycosylated PSMs results in higher scoring false matches, requiring a higher score threshold for effective filtering than for non-glycosylated PSMs. Since non-glycosylated PSMs make up the majority of the results, filtering all PSMs together would yield an insufficiently low score threshold for glycosylated PSMs. After the group-specific 1% FDR filter was applied to glycosylated and non-glycosylated PSMs, 1% peptide- and protein-level FDR filters were applied. A sequential filtering step was then applied to remove any PSMs matched to proteins that did not pass the 1% protein-level FDR. Glycan assignment was subsequently performed in PTM-Shepherd using the default N-glycan database^29^ and parameters along with a 0.05 glycan q-value threshold.

### Deconvolution of the MHC associated peptides

Motif deconvolution is the process of finding HLA-binding motifs and their corresponding binding core offsets for a set of peptides. A first deconvolution that required manual inspection was performed using MoDec^38^. The peptides were grouped by subject (*i*.*e*., instances of the same replicates). A maximum of 10 clusters, 20 runs, and a minimum peptide length of 12 amino acids were considered. Since HLA-II ligands from the same subject come from different alleles, MoDec provides a direct interpretation and assigns peptides with similar binding cores to clusters (i.e., HLA motifs). However, manual inspection is still required to (I) the number HLA motifs MoDec detected per subject and (II) annotate these motifs (*i*.*e*., clusters) to their respective HLA II alleles. Hence, the MoDec-identified HLA motifs were assigned to the correct HLA class II alleles by manual inspection for each analyzed subject. A second deconvolution that didn’t require manual inspection, inspired from Kaabinejadian *et al*. 2022^41^, was performed using NetMHCIIpan 4.1^42^. Briefly, all unique peptides were predicted for MHC presentation towards all the MHC alleles expressed in the given subject. The likelihood of peptides being presented by a given MHC molecule is given by the percentile rank score, which ranges from 0 to 100, with 0 being the strongest binding score. Peptides with a percentile rank score > 20 were considered MS co-immunoprecipitated contaminants and labeled as trash. Peptides with a percentile rank score ≤ 20 were assigned to the lowest scoring allele of a given subject. We applied the second deconvolution method using NetMHCIIpan 4.1 to the entirety of the subjects in this study, considering the similarity of the results to the first deconvolution method (*i*.*e*., MoDec).

### Figure generation

Motif plots were generated using the Python library Logomaker^52^, heatmaps using seaborn^53^ and other plots using matplotlib^54^.

## Authorship contribution

G.B. collected and curated the data, generated the figures and supplementary materials, and drafted and coordinated the manuscript. D.P. performed the immunopeptidomics analysis, supported figure generation, interpretation of results, and drafting and coordination of the manuscript. Y.H. produced the web portal and helped to revise the manuscript. F.Y. supported the study with software development related tasks. F.L. supported the study by adding a group-specific FDR feature to Philosopher. J.A. supported with the writing of the manuscript. M.C. helped with the study design and manuscript revision. A.I.N. conceived the project, helped with the study design and revision of the manuscript, and provided overall supervision.

## Acknowledgments

This work was funded in part by NIH grants R01-GM-094231, U24-CA210967, U24-CA271037, and by the International Centre for Cancer Vaccine Science, a project carried out within the International Research Agendas programme of the Foundation for Polish Science and co-financed by the European Union under the European Regional Development Fund.

