## Supplementary Figure 1 for "HLA-Glyco: A large-scale interrogation of the glycosylated immunopeptidome"

### Supplementary materials

#### Supplementary Figure 1

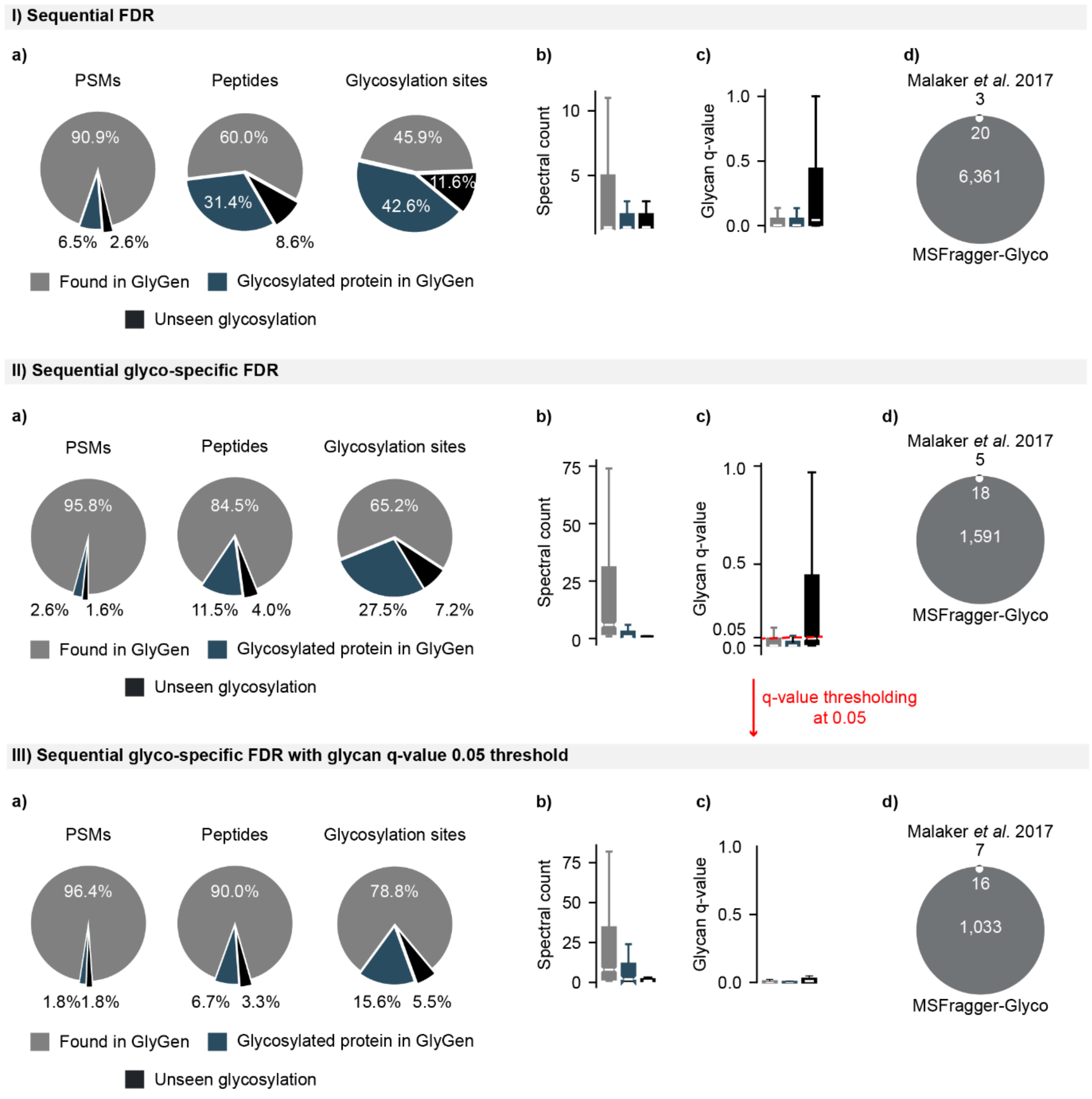


**Supplementary Figure 1: Comparison of 3 different FDR control strategies for HLA glycosylated peptides. Strategy I** referred to as “sequential FDR” is typically used with enzymatic (i.e., trypsin) glycoproteomic searches. **Strategy II** referred to as “sequential glyco-specific FDR” has been developed in this study to handle non-specific (*i.e.,* non-specific cleavage of proteins at every peptide bond) glyco searches. **Strategy III** is the one being used in this study and consists of applying the sequential glyco-specific FDR with an additional glycan q-value threshold of 0.05. **a)** Percentage of glyco-PSMs, glycopeptides and glycosylation sites found in GlyGen. Peptides with glycosylation sites reported in GlyGen are shown in gray, within glycosylated protein are shown in blue, and unreported are shown in black. **b)** Abundance of the 3 categories from panel (a) by spectral count. **c)** The glycan q-value range of the 3 categories from panel (a). **d)** Comparison of the identified glycosylation sites identified in this study with Malaker *et al.* 2017 findings.
